# Identification of VIMP as a gene inhibiting cytokine production in human CD4+ effector T cells

**DOI:** 10.1101/2020.07.06.189456

**Authors:** Christophe Capelle, Ni Zeng, Egle Danileviciute, Sabrina Freitas Rodrigues, Markus Ollert, Rudi Balling, Feng Q. He

## Abstract

Many players regulating the CD4^+^ T cell-mediated inflammatory response have already been identified. However, the critical nodes that constitute the regulatory and signalling networks underlying CD4 T cell responses are still missing. Using a correlation network-guided strategy based on time-series transcriptome data of human CD4^+^CD25^-^ effector T cells (Teffs), here we identified *VIMP* (VCP-interacting membrane protein), one of the 25 genes encoding selenocysteine in humans, as a gene regulating the effector functions of human CD4 T cells. Knocking-down *VIMP* in Teffs enhanced their proliferation and expression of several cytokines, including IL-2 and CSF2. We identified VIMP as an endogenous inhibitor of cytokine production in Teffs via both, the E2F5 transcription regulatory pathway and the Ca^2+^/NFATC2 signalling pathway. Our work not only indicates that VIMP might be a promising therapeutic target for various diseases involving CD4 T cells, but also shows that our network-guided approach might be generally applicable to different types of cells and can significantly aid in predicting new functions of the genes of interest.

**One-sentence summary:** Using a network-guided approach, we identified that Selenoprotein S (SELS or VIMP) negatively regulates cytokine expression of human CD4+ effector T cells via the E2F5 and calcium Ca2^+^/NFATC2 pathways.

## Introduction

CD4^+^ T cells represent a major subset of immune cells that are crucial for mounting and regulating an adequate immune response. However, during many acute infectious and complex chronic diseases, those T cells are dysregulated, either having an impaired responsive capacity or causing adverse effects through self-recognition and/or overactivation. Therefore, rebalancing the CD4^+^ T cell-mediated inflammatory response has been essential for the design of therapeutic options for those diseases (*1*). Although, many players regulating the inflammatory response, cytokine production and differentiation of CD4^+^ T cells have already been identified in the past (*2-5*), a thorough understanding of the regulatory and signaling networks governing inflammatory cytokine production in Teffs is still missing. The gap is not only attributable to the long-standing nature of traditional trial-and-error experimental procedures, but also due to the lack of the reliable high throughput computational prediction.

VIMP, also known as Selenoprotein S (SELS), SELENOS, TANIS or SEPS1, is one of the only 25 genes encoding the 21st amino acid selenocysteine in humans (*6*). Located in the endoplasmic reticulum (ER) membrane, VIMP is mainly known as an important component of the ER-associated degradation (ERAD) complex (*7, 8*) and physically binds to several ER membrane proteins (*9, 10*). VIMP plays a role in mediating retro-translocation of misfolded proteins from the ER lumen to the cytosol, where the ubiquitin-dependent proteasomal degradation takes place (*11*). Genome-wide association studies have shown that polymorphisms in the promoter region of *VIMP* are linked to a wide spectrum of common complex diseases, including cardiovascular disease (*12*), diabetes (*13, 14*), cancer (*15-17*), sepsis (*18*) and autoimmune diseases (*19, 20*), in which activation of the immune system is believed to be dysregulated (*21*).

Meanwhile, dysfunction of the ER and the unfolded protein response causes intestinal inflammatory diseases in several murine models (*22*). Additionally, a reduced expression of VIMP causes an increased expression of inflammatory cytokines, such as IL6, IL1β and TNFα in macrophages (*23*), as well as IL1β and IL6 expression in astrocytes (*24*). However, other studies did not show significant association between VIMP and the examined human inflammatory diseases (*25*). This controversy underlines the necessity for a better understanding of how VIMP contributes to the pathogenesis of some inflammatory diseases, i.e., through which cell types and which molecular pathways VIMP contributes to the observed dysregulated inflammatory responses. Therefore, we sought to investigate whether and how VIMP plays a role in relevant specific immune cells, e.g., CD4^+^ T cells, a key subset of immune cells orchestrating different types of immune responses and being heavily involved in different complex diseases.

We have previously developed a correlation-network guided approach, based on the guilt-by-association theory (*26-28*), to predict novel key genes of a given biological process or function and have successfully applied it to human CD4^+^CD25^high^CD127^low^ regulatory T cells (Tregs) (*29, 30*). Here, we extended the strategy to human CD4^+^ effector T cells (Teffs) that were derived and expanded from sorted CD4^+^CD25^-^ T cells by co-culturing with EBV-transformed B cells and were able to predict that VIMP might play an important role in regulating the effector responses of Teffs. Combining both the network analysis and experimental verification, we identify *VIMP* as a previously unreported vital endogenous inhibitor of cytokine production in CD4^+^ Teffs and reveal the molecular mechanisms through which VIMP regulates CD4^+^ Teff responses.

## Results

### VIMP is temporally upregulated after TCR stimulation in Teffs

Using our previously reported high-time-resolution time-series (HTR) transcriptome data of Tregs and Teffs following TCR (T cell receptor) stimulation in the first six hours (*29*), we observed that in Teffs the transcript level of VIMP temporally peaked within 2-3 hours following stimulation, which was followed by a gradual decrease (**Fig. 1A**). In contrast, the *VIMP* mRNA level was kept almost constant in Tregs during the first six hours following TCR stimulation (**Fig. 1A**), indicating a possible specific role for VIMP in Teffs. Our quantitative real-time PCR (qPCR) results validated the transitionally elevated expression of the VIMP transcript in Teffs isolated from different healthy donors (**Fig. 1B**). We also observed a correlation over time between the transcription levels of *VIMP, IL2, IL13* and *CSF2* following TCR stimulation, indicating a potential regulatory effect between *VIMP* and some of the cytokines in Teffs (**Fig. 1B**). By flow cytometry (**Fig. 1C**), we confirmed the gradual upregulation of VIMP protein expression in the first 5 hours following TCR stimulation. In summary, both mRNA and protein expression of VIMP were upregulated following TCR stimulation, which was correlated with the expression of several examined cytokines, indicating a potential role of VIMP in regulating Teff responses.

**Figure 1.**
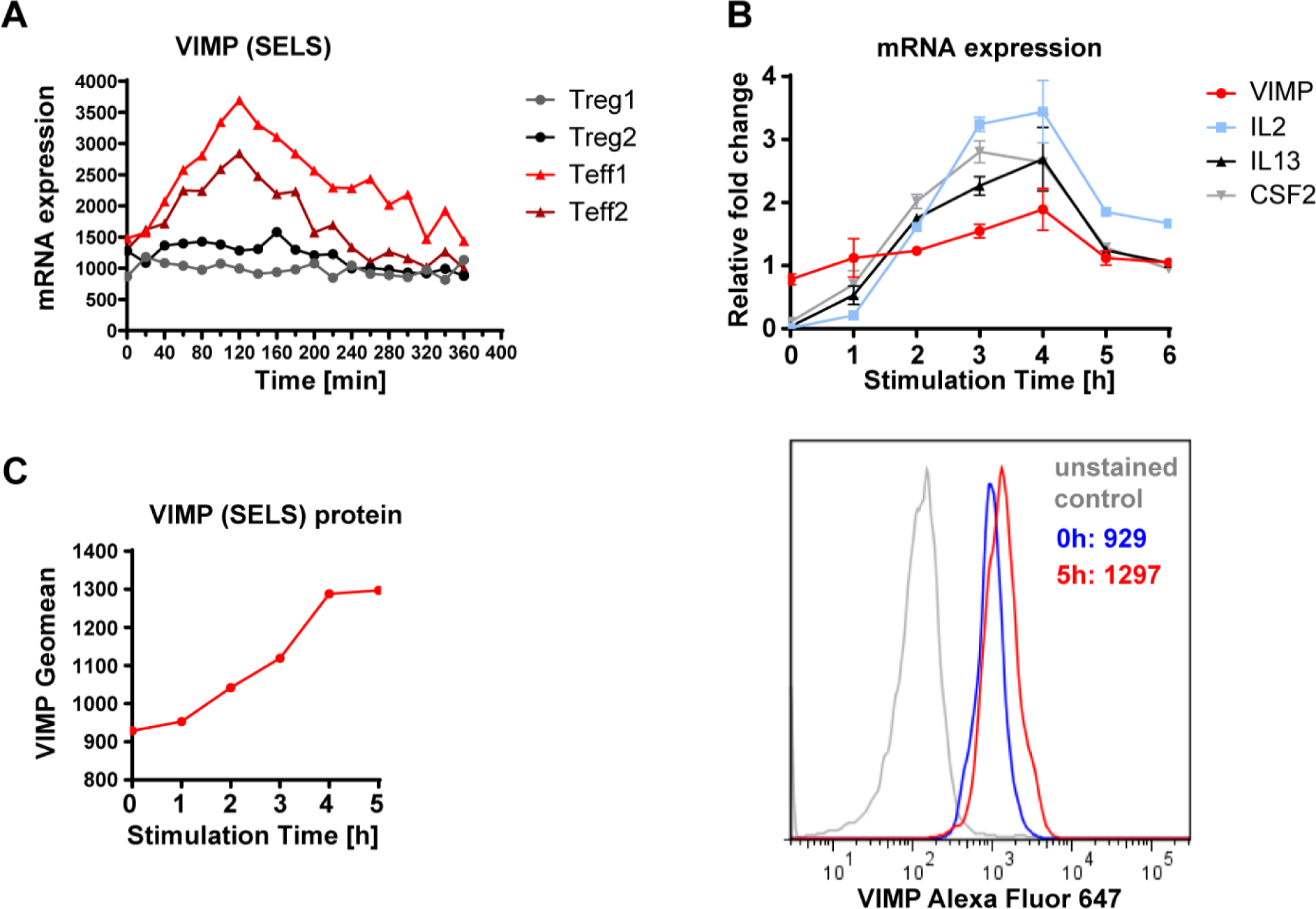
VIMP is temporally upregulated in Teffs following TCR stimulation. (**A**) The kinetics of transcriptional expression of VIMP in the first 6 h following anti-CD3/-CD28 stimulation assessed by HTR time-series microarray data. Teff1 and Teff2 are the two independent repeated HTR time-series experiments from different donors. (**B**) Representative experiments, reproduced in 4 independent donors showing mRNA expression of *VIMP, IL2, IL13* and *CSF2* measured by qPCR in Teffs stimulated by anti-CD3/-CD28 beads. The data represents the mean gene expression normalized to *RPS9*. Error bars represent standard deviation (SD) values. (**C)** Representative flow cytometry quantitative analysis showing elevation of VIMP protein expression in Teffs following TCR stimulation. Results represent four (**B**) and three (**C**) independent experiments of different donors.

### VIMP inhibition upregulates cytokine expression of Teffs

The upregulation of VIMP and its correlation to cytokine expression encouraged us to further investigate VIMP’s potential role in CD4 T cell responses. We and others have previously shown that the enriched pathways or processes or functions among the genes surrounding a given gene in the correlation network might give valuable indications on potential new functions of the given hub gene (*29, 30*). Therefore, we used our correlation network-guided approach to predict the potential functions of VIMP by identifying the enriched pathways among the genes that are linked to *VIMP* within the subnetwork of the Teff correlation network, which was extracted from our published HTR datasets and networks (*29*) (**Fig. 2A**).

**Figure 2.**
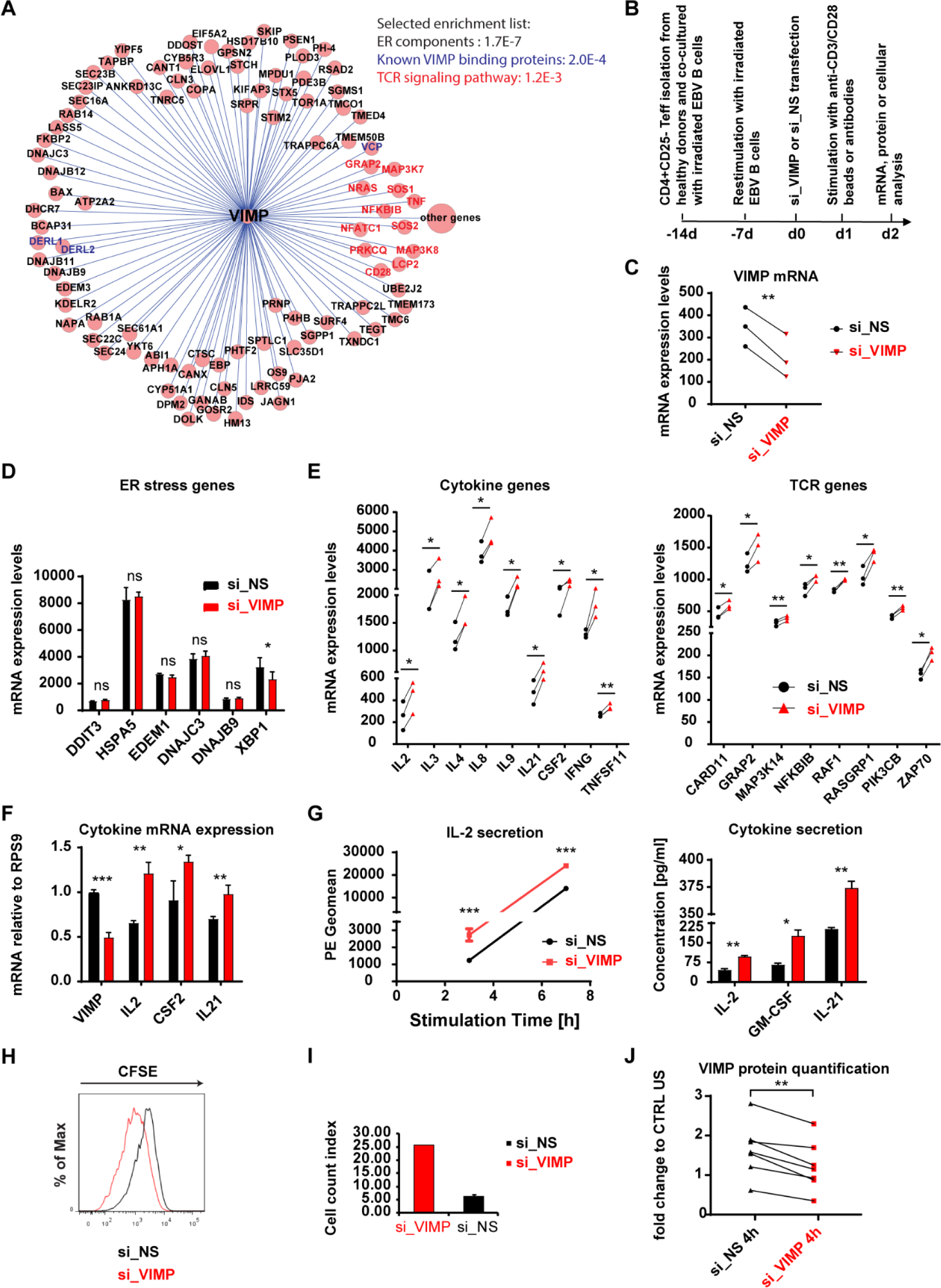
VIMP controls cytokine expression in Teffs and interferes with the TCR signalling pathway. (**A**) The *VIMP* subnetwork extracted from the constructed Teff correlation network based on the HTR transcription microarray data. Each circle represents one gene. Each line between *VIMP* and the other genes represents a correlation link. The selected list of significantly enriched pathways or components is displayed (the P-value resulted by cumulative Binomial distribution test was provided for each item). (**B**) Schematic of the experimental flow for the stimulation, gene silencing and analysis of Teffs. (**C)** mRNA expression showing the significant knockdown of VIMP in the microarray experiment. **(D, E**) Microarray data showing the fold change or expression values in the mRNA expression of ER-stress responsive genes (**D**), cytokine and TCR signaling genes (**E**). We only presented the transcripts with P-values lower than 0.05 by both PLIER and RMA methods and at least a 1.2 fold change in all 3 independent donors (for details, see Methods). (**F**) mRNA expression measured by qPCR of the genes *VIMP, IL2, CSF2* and *IL21* of Teffs following TCR stimulation and knockdown with either control non-specific scrambled siRNA (si_NS) or *VIMP*-specific siRNA (si_VIMP). Before stimulation, the cells were first transfected with siRNA for 1 day (for all the figures). (**G**) The concentration of the cytokines IL2, GM-CSF and IL21 detected in the cell culture medium following anti-CD3/-CD28 stimulation for different time points (left panel, IL2 alone by CBA measurement) or 8 hrs (right panel, multiplexing by MSD). PE geometric mean (Geomean) corresponds to the IL2 concentration signal in the media. (**H, I**) Proliferation of Teffs following TCR stimulation and VIMP knockdown, measured by CFSE proliferation assay (**H**) by counting the T cells following stimulation (**I**). Before Teffs were co-cultured with EBV-transformed B cells for 2 days, they were first transfected with siRNA for 1 day. (**J**) Quantification of the western blot protein bands and normalization of VIMP to the housekeeping gene GAPDH. Each dot represents one sample. CTRL US, unstimulated Teffs treated with si_NS. Data are mean± standard deviation (s.d.). The P-values are determined by a two-tailed paired Student’s t test (except for **A** and **G**). The results in **G** was analyzed using non-paired t test. ns or unlabeled, non-significant; *P<=0.05, **P<=0.01 and ***P<=0.001. Results represent three (**C-E**) and six (**F-J**) independent experiments of different donors.

Consistent with its known function and its localization in the ER membrane, the genes surrounding *VIMP* in the correlation network were significantly enriched for ER components (P=1.7E-7, cumulative Binomial distribution) (**Fig. 2A**). Furthermore, 3 out of the 10 experimentally-validated VIMP-binding partners found in the literature in other cellular types are directly linked to *VIMP* in the Teff correlation network (P=2.0E-4, http://string-db.org (*31*), indicating the reliability of our method. Surprisingly, the pathway enrichment analysis shows that the genes linked to *VIMP* are significantly enriched for components involved in the TCR signaling pathway (P=1.2E-3, cumulative Binomial distribution) (**Fig. 2A**), suggesting a potential role of VIMP in the Teff response according to our network-based analysis strategy (*29, 30*). However, the genes linked to the hub gene of interests in the correlation network could follow at least two scenarios (*32-34*). First, those genes could simply be co-regulated with the hub gene, but perform independent functions. Second, those genes could be co-expressed with the hub gene and play related roles in the same pathways to coordinate cellular resources for a particular function or purpose under certain conditions. We will test these possibilities in this work.

In order to systematically assess whether and how *VIMP* controls the inflammatory response of Teffs, we performed a transcriptome analysis of CD4 Teffs isolated from peripheral blood mononuclear cells (PBMC) of three healthy donors that were subjected to a specific-siRNA knockdown of VIMP (si_VIMP) or a control unspecific siRNA (si_NS) followed by anti-CD3/-CD28 stimulation (**Fig. 2B**). As shown in **Fig. 2C**, the mRNA expression of *VIMP* was significantly downregulated in Teffs by using siRNA knockdown.

As VIMP has reported functions in ER stress, we first checked the ER-stress responsive genes in the transcriptomic data of the Teffs transfected with si_VIMP vs. that treated with control siRNA (si_NS). By perturbing the expression of *VIMP*, we expected a change in expression of some ER-stress responsive genes. Nonetheless, our transcriptome data of Teffs with VIMP partial knockdown did not show any significant change in mRNA expression of those genes (e.g., *CHOP (DDIT3), GRP78 (HSPA5), EDEM1, DNAJC3 (P58IPK)* and *DNAJB9 (ERdj4)* (*35*)) (**Fig. 2D**). Only the expression of the ER-stress regulator XBP1 (*36*) was significantly but modestly decreased. Indeed, data from intestinal epithelial cells show that VIMP is only a marker, but not a regulator ER-stress (*37*). This shows that VIMP’s direct involvement in ER stress might not be ubiquitous to all cell types. We therefore ruled out the possibility that VIMP directly regulates the expression of the ER-stress responsive genes, indicating other roles of VIMP in modulating the Teff responses.

Considering that the TCR signaling pathways were significantly enriched in the VIMP correlation network, we further analyzed the genes related to the TCR signaling pathway in Teffs after *VIMP* knockdown. Notably, we found 13 significantly upregulated genes involved in the TCR signaling (refer to https://www.genome.jp/kegg-bin/show_pathway?hsa04660) in the microarray datasets of the Teffs, although subjected to only a partial knockdown of *VIMP*, including several cytokines, namely *IL2, IL4, CSF2* and *IFNG* (**Fig. 2E**). Moreover, transcripts of the key TCR related signaling molecules, such as *GRAP2, ZAP70, RASGRP1* and *RAF1* were significantly affected (**Fig. 2E**). With the observation in mind that *VIMP* and the TCR signaling related genes were directly linked in our HTR correlation network (**Fig.2A**), this effect of the siRNA perturbation was not unexpected. Our results suggest that VIMP negatively regulates the expression of specific cytokines and influences the expression of important components of the TCR signaling pathway.

To further confirm whether *VIMP* regulates cytokine expression in Teffs, using PBMC of independent donors we measured the cytokine mRNA expression by qPCR and the secreted cytokines of Teffs that were subjected to a *VIMP* knockdown. Indeed, *IL2, IL21 and CSF2* mRNA were significantly upregulated in stimulated Teffs transfected with si_VIMP, compared with control Teffs (with si_NS) (**Fig. 2F**). This observation was further consolidated by increased IL2, IL21 and GM-CSF protein production in the culture media of stimulated Teffs transfected with si_VIMP, compared with that treated with control scrambled siRNA (**Fig. 2G**). Furthermore, the VIMP knockdown also significantly promoted T cell proliferation as indicated by both CFSE dilution and Teff cell number counting experiments (**Fig. 2I**). As IL2 concentration was already significantly higher at 3 hours following stimulation (**Fig. 2G**) and no cell division was already expected, the enhanced IL2 secretion following VIMP knockdown was not simply caused by more Teffs. All the analyses were done under the precondition that VIMP protein was successfully silenced (**Fig. 2H**). In short, VIMP negatively regulates the expression of several cytokines in Teffs following stimulation.

### VIMP controls cytokine expression via the transcription factor E2F5

Next, we aimed to identify any (co-)transcription factors (TFs), whose expression were significantly affected after silencing VIMP, as they often serve as the key components orchestrating the activity of the relevant pathways. Through a systematic analysis of all the known mammalian TFs or co-factors (*38*) in our microarray datasets, *E2F5* was found as the most significantly upregulated TF, following a partial VIMP knockdown (**Fig. 3A**). Conversely, *RNF14* (ring finger protein 14) was the most downregulated cofactor together with the downregulated TFs such as *CEBPG* (CCAAT Enhancer Binding Protein Gamma*), ZBTB20* (zinc finger and BTB domain containing 20) and *IRX3* (Iroquois homeobox 3) (**Fig. 3A**). We further confirmed the expression change of these (co-)TFs by qPCR in independent healthy donors (**Fig. 3B**).

**Figure 3.**
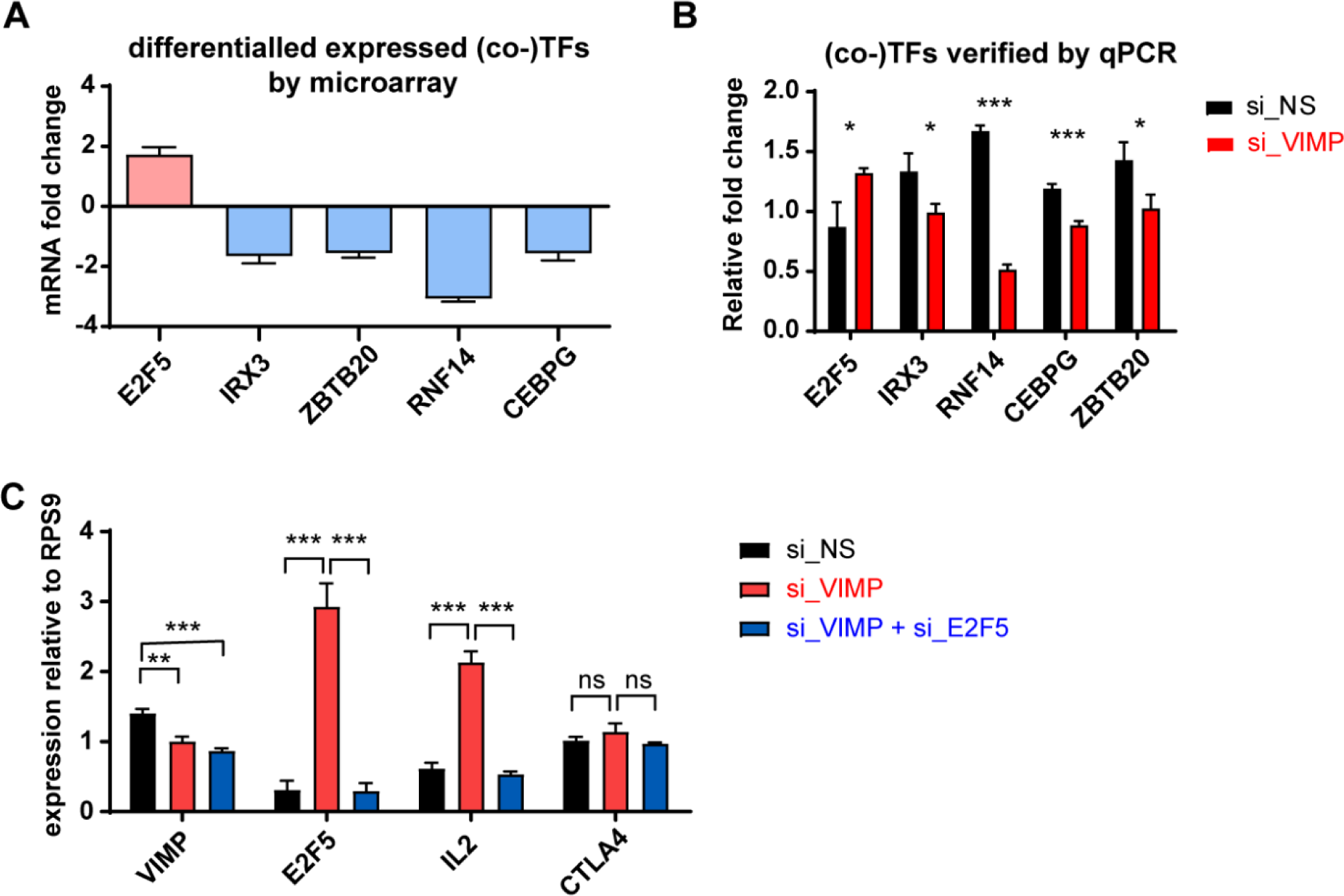
VIMP inhibits E2F5 to regulate IL2 expression. (**A**) The most significantly affected (co-)transcription factors selected from our microarray analysis; the y-axis indicates the fold change between Teffs transfected with siRNA specific against VIMP(si_VIMP) or non-specific scrambled si_RNA (si_NS). (**B**) mRNA expression measured by qPCR from Teffs of independent healthy donors of the genes displayed in **A** to confirm the change in their expression levels following *VIMP* knockdown. (**C**) mRNA expression measured by qPCR of the genes *VIMP, E2F5, IL2* and *CTLA4* of Teffs transfected with si_NS, si_VIMP, or both si_VIMP and si_E2F5. Data are mean± standard deviation (s.d.). The P-values are determined by a two-tailed paired Student’s t test. ns or unlabeled, non-significant; *P<=0.05, **P<=0.01 and ***P<=0.001. Results represent three (**A**) and four (**B-C**) independent experiments of different donors.

E2F5 has previously been reported to be a downstream target of IL-2 in an immortalized human T cell line (*39*). But to our knowledge, there are no reports yet of E2F5 sitting at the upstream pathway regulating inflammatory responses, especially cytokine production. Nevertheless, being the most significantly upregulated TF following a partial knockdown of VIMP, we assumed that E2F5 might be an important component in the regulatory pathway through which VIMP regulates the Teff inflammatory response.

Therefore, we decided to investigate whether VIMP controls the cytokine expression by negatively regulating E2F5 expression in stimulated Teffs. In order to examine this hypothesis, we silenced *VIMP* alone or in combination with *E2F5* and measured the expression of selected cytokines by qPCR. In addition to the reduced expression of *VIMP*, the upregulation of *E2F5* expression that was driven by VIMP knockdown was abolished in the *VIMP* and *E2F5* double knockdown Teffs (**Fig. 3C**). While silencing *VIMP* alone upregulated *IL2* expression in stimulated Teffs, a dual knockdown of *VIMP* and *E2F5* suppressed the surge of *IL2* caused by VIMP knockdown alone (**Fig. 3C**). Even though E2F5 is a general regulator of transcription, we did not observe any effect of *E2F5* knockdown on genes that are not directly involved in Teff inflammatory response, such as *CTLA4* **Fig. 3C**). This excluded a generalized effect of E2F5 on the transcription regulation in Teffs. In brief, our data support that VIMP regulates the expression of inflammatory cytokines, i.e., IL2, by restraining the expression of the TF E2F5 in Teffs.

### VIMP controls cytokine expression via the Ca^2+^/NFATC2 signaling pathway

To further delineate VIMP’s regulatory pathways beyond the altered expression of individual TFs determined by the differential-expression analysis of our microarray datasets, by applying the Ingenuity Pathway Analysis (IPA) we mapped the up- or down-regulated cytokine and TCR related genes into the known regulatory network structures. We found that many of those differentially-expressed genes are controlled by the expression change of the so-called hub genes *IL2, RAF1, IL21, TNFSF11*, as well as nuclear factor of activated T cells (NFAT) activity (**Fig. 4A**). Although *NFAT* transcript expression was not significantly affected (**Fig. 4A**), its activity was predicted to be increased by the IPA computational analysis. Meanwhile, we investigated the VIMP subnetwork in the Teff correlation network in more depth (**Fig. 2A**). We found that genes for several components of NFκB, NFAT and MAPK signaling pathways were also directly linked to *VIMP*, indicating that those pathways might be involved in the regulation of the inflammatory response of Teffs by VIMP. In order to determine whether any of the relevant signaling pathways downstream of the TCR pathway that were suggested by the computational analysis are affected by VIMP expression, we quantitatively assessed the phosphorylation levels of up to 10 various signaling proteins by flow cytometry (**Fig. 4B**). Canonical (NFKB1 p105 and p65) and non-canonical (NFKB2 p100 and RELB) NFκB signaling pathways, as well as several MAP kinase sub-pathways (ERK1/2, p38, JNK1/2 and cJun) were not significantly affected in their phosphorylation levels (**Suppl. Fig. 1A-E, 2**). The phosphorylation level in one of the NFAT family members, NFATC1, was also not significantly affected by VIMP knockdown in stimulated Teffs (**Suppl. Fig. 1F, 2**). However, the phosphorylation level at the specific site Ser326 of another NFAT family member, NFATC2 (also known as NFAT1) was significantly reduced even following a partial VIMP knockdown, as quantified by both flow cytometry and western blotting in Teffs isolated from different donors (**Fig. 4C-E, Suppl. Fig.1G**). Total NFAT protein remained unaffected by the partial VIMP knockdown (**Suppl. Fig. 1H**). In resting T cells, NFAT proteins are phosphorylated and reside in the cytoplasm (*40, 41*). In order to be able to translocate to the nucleus and induce gene expression, NFAT is de-phosphorylated following the TCR signaling. As the NFAT activity is known to regulate IL2 expression in T cells (*42*), the observed downregulation of NFATC2 phosphorylation, following *VIMP* knockdown, showed that the upregulation of IL2 expression was, at least in part, due to an increase in NFAT activity.

**Figure 4.**
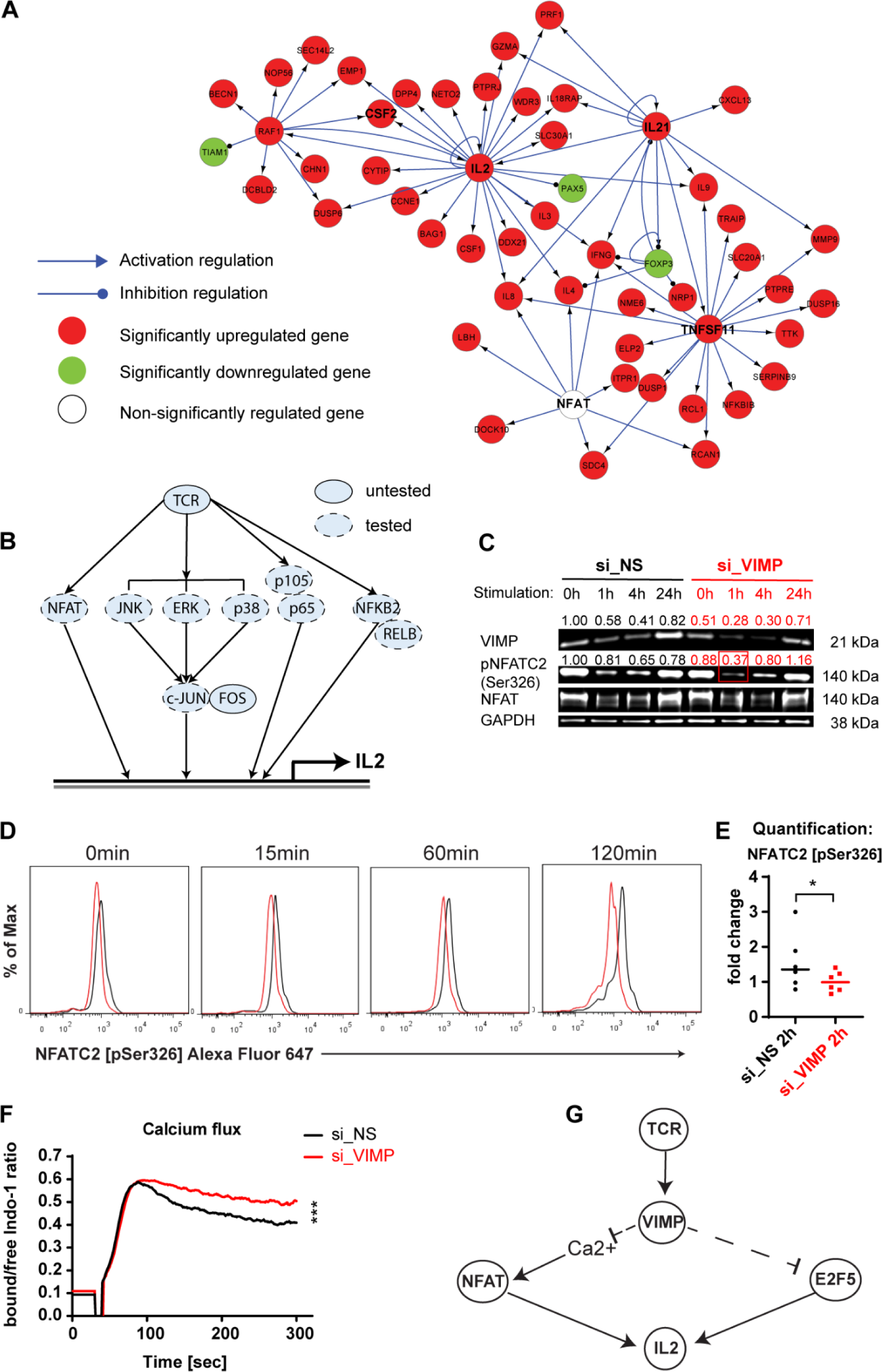
VIMP controls cytokine expression via the Ca2+/NFATC2 phosphorylation pathway. (**A**) Network representation of the cytokine and TCR related genes affected by the knockdown of VIMP by Ingenuity Pathway Analysis (IPA). Red, significantly upregulated genes; green, significantly downregulated genes; white, non-significantly affected gene at the transcriptional level. The link with arrow indicates a known direct or indirect positive transcription regulation, the link with circle indicating a negative one from the IPA knowledge databases. (**B**) Graphical representation of the major signaling pathways downstream of the TCR signaling, (un)tested for their phosphorylation levels. (**C, D**) Phosphorylation of NFATC2 (NFAT1) in Teffs assessed by western blot (**C**) or flow cytometry (**D**) at different time points following anti-CD3/-CD28 stimulation (**C**) or PMA/Ionomycin stimulation (**D**). Before stimulation, Teffs were first transfected with specific siRNA against VIMP (si_VIMP) or non-specific siRNA (si_NS) for 24 hrs. (**D**) Representative flow-cytometry plots of pNFATC2 in Teffs. (**E**) Pooled pNFATC2 data from multiple donors at 120 mins post stimulation. For **D** and **E**, only gated viable Teffs were displayed for all the phosphorylation results. The y-axis represents the percentage of maximum (scales each curve to mode=100%) (% Max). The fold change was calculated by normalizing the geometric mean (Geomean) of the fluorescence intensities of all the conditions to that of the unstimulated control siRNA knockdown condition. (**F**) Representative graph out of 3 independent experiments for the calcium flux in Teffs following stimulation. The one displayed here used Ionomycin stimulation after si_VIMP or si_NS transfection for 24h. The y-axis represents the ratio between calcium bound and free Indo-1 dye over time. (**G**) Graphical representation summarizing the two mechanisms through which VIMP regulates cytokine expression in CD4 Teffs. The P-values are determined by a two-tailed paired Student’s t test. ns or unlabeled, non-significant; *P<=0.05, **P<=0.01 and ***P<=0.001. Results represent three (**C, F**), six (**D, E**) independent experiments of different donors.

The distinguishable feature of NFAT is that it relies on Ca^2+^ influx and subsequent Ca^2+^/calmodulin-dependent phosphatase calcineurin to become dephosphorylated and being able to translocate to the nucleus to induce gene expression (*43*). Although *VIMP* has not yet been linked to the calcium signaling, other selenoproteins have been described to regulate the calcium signaling and homeostasis (*44*). We therefore further asked whether *VIMP* knockdown affects the calcium flux in Teffs and measured it by flow cytometry using the calcium indicator Indo-1. Indeed, the Teffs in which *VIMP* was silenced vs. the control Teffs, showed a significantly higher flux of Ca^2+^ ions (**Fig. 4F**), further illustrating the increased NFATC2 activity and IL2 expression.

In summary, our data strongly suggest that *VIMP* inhibition upregulates the expression of cytokines, such as IL2 by two mechanisms at different levels (**Fig. 4G**). On the transcription regulatory level, *VIMP* controls the expression of transcription factor E2F5 and multiple genes involved in the TCR signaling and the inflammatory response. On the signaling transduction level, *VIMP* knockdown modulates Teff responses by controlling the Ca^2+^ flux and the downstream NFATC2 de-phosphorylation.

## Discussion

So far many important components in the regulatory or signaling networks modulating the inflammatory responses of Teffs still remain elusive. With the development of systems biology, researchers have greater opportunities to use top-down approaches to objectively infer and identify novel key genes or proteins in the process of interest.

In this work, we have applied our previously published correlation network-guided strategy to predict new genes regulating the effector functions of CD4^+^CD25^-^ Teff cells, i.e., cytokine production. We identified VIMP, an ER membrane protein encoding the 21^st^ amino acid selenocysteine, as a novel negative regulatory gene of the Teff response. By thoroughly analyzing our transcriptomic datasets and experimentally verifying the most promising candidates, we were able to identify the Ca^2+^/NFATC2 signaling pathway as the primary pathway through which VIMP inhibition enhances cytokine production of Teffs. Coincidently, the others have recently reported a co-expression of VIMP and NFATC2 transcripts within the murine interfollicular epidermis using single-cell RNA-seq analysis, from another angle indicating that our conclusion might hold true (*45*). We have also shown that E2F5 plays a significant role in the VIMP-mediated regulation of the Teff IL2 expression. However, whether the E2F5 pathway and the Ca2+/NFATC2 signaling pathway controls VIMP-mediated IL2 expression in a sequential manner or in parallel requires further investigation.

In our transcription-factor (TF) focused analysis, we identified not only E2F5, as the most upregulated TF, but also several downregulated TF genes, following VIMP knockdown. Among those downregulated ones, RNF14 (ring finger protein 14), a less characterized gene, represented the most significantly downregulated co-factor, attributable to VIMP knockdown in Teffs. Although very limited, a published report shows that RNF14 modulates the expression of inflammatory and mitochondria-related genes in a murine myoblast cell line (*46*). Another downregulated TF *ZBTB20* has originally been studied in human dendritic cells (*47*), later in myeloid cells (*48*) and B cells (*49*) to regulate their effector functions and differentiation. The Iroquois homeobox 3 (*IRX3)* has been recently linked to CD8 T cell survival and fate determination in one preprint (*50*). Although there is no evidence of CEBPG being involved in the regulation of cytokine expression in CD4 effector T cells, other C/EBP protein family members were shown to act as negative regulators in the production of inflammatory cytokines (*51, 52*). Therefore, those TFs might deserve a further investigation.

At first glance, those genes as well as NFAT and E2F5, do not show a high overlap in their known protein networks (http://string-db.org) (*31*), indicating that those pathways might represent independent axes. A closer look, however, shows that RNF14 is a positive regulator of WNT signaling by binding TCF/LEF TFs in colon cancer cells (*53*). Conversely, NFAT signaling was reported to be antagonistic to WNT signaling in neural progenitor cells, thus promoting their differentiation (*54*). Our results showed a decrease in RNF14 expression and an increase in NFAT activity following VIMP knockdown in Teffs. In line with their involvement in WNT signaling in other cell types, the significant change in expression or activity of RNF14 and NFAT could synergistically contribute to the inhibition of WNT signaling in Teffs, consequently impairing T cell function and development (*55*). These reports and our observation suggest RNF14 another promising candidate meriting further investigation to better understand the regulatory function of VIMP in the Teff inflammatory response.

Overall, using both hypothesis-free top-down computational analyses and bottom-up experimental methods, we have shown a regulatory role for the selenoprotein VIMP in controlling cytokine expression in CD4^+^CD25^-^ Teffs by affecting several signaling pathways and transcriptional regulatory pathways. The same strategy should be generally extendable to other cell types in assisting the prediction and discovery of novel functions of other genes of importance, which is one of the essential aims for a wide spectrum of various researchers in biomedicine.

Selenoproteins fully rely on selenium for their biosynthesis and function. Dietary selenium supplementation in mice has been shown to increase the biosynthesis of VIMP (*56, 57*) and to reduce the expression of several inflammatory cytokines, such as tumor necrosis factor α (TNFα), monocyte chemoattractant protein 1 (MCP1) and IL2 (*56, 58-60*). Dietary selenium supplementation has further been linked to alleviate several complex and multifactorial diseases (*61-64*). On the other hand, selenium deficiency in mice results in an increased pathology from influenza viral infections, due to an exaggerated inflammatory immune response (*60*). In summary, our data identified an unrecognized critical regulatory role of the selenoprotein VIMP in the inflammatory responses of human CD4^+^ Teffs. Our observation provides a novel viable insight into how dietary supplementation of selenium might mediate its effects in CD4^+^ Teffs and underscores the potential in therapeutically targeting VIMP in the treatment of various inflammatory and inflammation-related diseases.

## Materials and Methods

### Primary T cell isolation and culture

Buffy coats from healthy donors were provided by the Red Cross Luxembourg and the informed consent was obtained from each donor by the Red Cross Luxembourg. The RosetteSep™ Human CD4+ T cell Enrichment Cocktail (15062, Stemcell) was added to undiluted blood at a concentration of 50 µl/ml and incubated for 30 min at 4°C. The blood was then diluted 2 times with FACS buffer (PBS + 2% FBS) and the CD4+ cells were isolated by gradient centrifugation at 1200 g for 20 min, using Lympoprep (07801, StemCell) and SepMate™-50 tubes (85450, Stemcell). CD4+ cells were stained with mouse monoclonal [RPA-T4] anti-human CD4 FITC (555346, BD Biosciences) (dilution 1:20), mouse monoclonal [M-A251] anti-human CD25 APC (555434, BD Biosciences) (dilution 1:20) and LIVE/DEAD® Fixable Near-IR Dead Cell Stain (L10119, ThermoFisher Scientific) (dilution 1:500). Primary CD4 T cells (CD4^+^CD25^-^) were then sorted on a BD Aria III Flow cytometry cell sorter (BD Biosciences).

| Target | Fluorochromes | Dilution | Company | Clone | Reference |
| --- | --- | --- | --- | --- | --- |
| CD4 | FITC | 1:20 | BD Biosciences | RPA-T4 | 555346 |
| CD25 | APC | 1:20 | BD Biosciences | M-A251 | 555434 |
| Live/Dead | Near Infra-Red | 1:500 | ThermoFisher Scientific | N.A. | L10119 |

Sorted CD4+ were cultured in IMDM (21980-032, ThermoFisher Scientific) complete medium, supplemented with 10% heat-inactivated (56°C, 45 min) fetal bovine serum (FBS) (10500-064, ThermoFisher Scientific), 1x Penicillin+Streptomycin (15070-063, ThermoFisher Scientific), 1x MEM non-essential amino acids (M7145, Sigma-Aldrich) and 1x β-mercaptoethanol (21985-023, ThermoFisher Scientific). Every seven days for a maximum of four weeks, Teffs were derived from isolated CD4^+^CD25^-^ T cells by restimulating them with irradiated Epstein–Barr virus (EBV) transformed B-cells (EBV-B cells) (*65*), at a 1:1 ratio to expand and maintain the culture. The EBV-B cells were irradiated in RS2000 X-Ray Biological Irradiator (Rad Source Technologies) for 30 min with a total of 90 Gy.

### Teff siRNA knockdown and stimulation

Targeted gene’s expression was knocked-down in up to 5 x 10^6^ cells using the P3 Primary Cell 4D-Nucleofector X Kit L (V4XP-3024, Lonza) with 90 µl P3 Primary cell solution and 100 pmol of corresponding si_RNA (resuspended in 10 ul RNAse-free H2O): si_Non-Specific scrambled control siRNA (si_NS or si_CTRL) (SC-37007, Santa Cruz), si_VIMP/SELS (SI03053512, Qiagen), si_E2F5 (SI00030436, Qiagen). The Amaxa 4D-Nucleofector™ X System (Lonza) was used to perform the electroporation and siRNA transfection according to the manufacturer’s recommended program for primary human T cells (with the program code EO-115). After transfection, the Teffs were transferred into a 12-well plate with pre-warmed complete medium and kept at 37 °C for 24 hours before being stimulated with 25 µl/ml of soluble antibodies (Immunocult™ Human CD3/CD28 T Cell Activator) (10971, StemCell), or 10ng/ml PMA (Phorbol 12-myristate 13-acetate, P8139, Sigma-Aldrich) and 100 ng/ml Ionomycin (I0634, Sigma-Aldrich) or Dynabeads® Human T-Activator CD3/CD28 for T Cell Expansion and Activation (11131D, ThermoFischer Scientific) (with 1:1 ratio between number of cells and beads) in a 24-well plate for different specified time periods.

### RNA extraction

RNA was extracted using the RNeasy Mini Kit (74106, Qiagen), following the manufacturer’s instructions and including the digestion of genomic DNA with DNAse I (79254, Qiagen). The cells were lysed in RLT buffer (Qiagen), supplemented with 1% beta-Mercaptoethanol (63689, Sigma-Aldrich). RNA concentration was measured by NanoDrop 2000c Spectrophotometer (ThermoFisher Scientific). For the microarray analysis, the quality of RNA was first checked by assessing the RNA integrity number (RIN) using the Agilent RNA 6000 Nano kit (5067-1511, Agilent) and the Agilent 2100 Bioanalyzer Automated Analysis System (Agilent), according to the manufacture’s protocol. Only the samples with RIN of 8.5 or higher were considered for further analysis.

### Microarray measurement and analysis

The transcriptomic analysis of human effector T cells expanded from CD4+CD25-T cells isolated from the PBMCs of healthy donors were performed on the Affymetrix human gene 2.0 ST array at EMBL Genomics core facilities (Heidelberg, led by Dr. Benes Vladimir). The facility used 500 ng of total RNA in the protocol with the Ambion® WT Expression Kit (cat. 4411974) in order to obtain 10 ug of cRNA, which was then converted to ssDNA. 5.5 ug of ssDNA was labeled and fragmented using the WT Terminal Labeling, polyA and hyb Controls Kit (Affymetrix, cat. 901524). 3.75 ug of fragmented/labeled ssDNA (with hybridization controls) was hybridized to Affymetrix HuGene 2.0 Genechip at 45 °C for 16 h with rotation (60rpm) and washed and stained on GeneChip Fluidics Stations 450 using GeneChip® Hybridization Wash and Stain Kit (Affymetrix, cat. 900720). Arrays were scanned using GeneChip Scanner 3000 7G with GeneChip Command Console software.

The expression signal at the exon level was summarized by the Affymetrix PLIER approach using the sketch approximation of quantile normalization with the option PM-GCBG (a GC content based background correction) using Affymetrix Expression Console v1.3.1.187. Before performing differential analysis, we first pre-processed the data with certain filtering steps. The filtering steps following the PLIER summary method included: 1) first removing any probeset whose cross-hyb type was not equal to 1; 2) removing any probeset corresponding to no identified gene or multiple genes according to the annotation (the file HuGene-2_0-st-v1.na33.2.hg19.transcript) and the library version r4 (May 23, 2012); 3), excluding the probesets with the average expression value in both groups (si_NS and si_VIMP) ≤ 2 times of the median value of the arrays (in our case, 2x the median was equal to the intensity value of 170); 4) if the mean intensity of the probesets in one group was higher, the number of absent calls among the three biological replicates should not be ≥ 1 in the group with higher mean intensity. To secure more robust analysis, we also analyzed the dataset using another model-based method (*66, 67*), i.e., RMA-sketch summary/normalization method (of note, the filtering steps mentioned above did not apply to the data resulted by the RMA-sketch summary method). We selected the probeset for further analysis only if the two-sided pair-wised T-test generated a P-value lower than 0.05 from the datasets summarized by both PLIER and RMA methods as demonstrated somewhere else (*67*). To obtain a certain number of starting candidates, we lowered the threshold of the change fold to 1.2, which had to be recurrent in all the three donors, for our further analysis in consideration of both facts that VIMP is not a (co)transcription factor and the siRNA knockdown efficiency was not 100%. The database of mammalian transcription factors or cofactors, or chromatin remodeling factors was downloaded from the work of others (*38*).

In this way, around 800 genes were significantly upregulated and around 550 genes were downregulated following VIMP knockdown, which were used for further analysis.

### Correlation network and IPA

The Teff correlation network based on high-resolution time series datasets of Teffs was already calculated and constructed in our previous work (*29*) and we extracted the VIMP subnetwork for a deeper analysis in this work. Ingenuity Pathway Analysis (IPA) was used to reconstruct the regulatory network from the Ingenuity database following the instruction of provider (QIAGEN).

### cDNA synthesis

A maximum of 500ng of RNA was used for human cDNA synthesis, using the SuperScript™ IV First Strand Synthesis System (18091050, ThermoFisher Scientific) and following the manufacturer’s instructions. The mastermix for the first step contained per sample: 0.5 µl of 50 µM Oligo(dT)20 primers (18418020, ThermoFisher Scientific), 0.5 µl of 0.09 U/µl Random Primers (48190011, ThermoFisher Scientific), 1 µl of 10 mM dNTP mix (18427013, ThermoFisher Scientific) and RNAse free water for a final volume of 13 µl in 0.2 ml PCR Tube Strips (732-0098, Eppendorf). The reaction tubes were transferred into a C1000 Touch Thermal Cycler (Bio-Rad) or UNO96 HPL Thermal Cycler (VVR) and subjected to the following program: 5 min at 65°C, followed by 2 min at 4 °C. For the second step, the reaction mix was supplemented with 40 U RNaseOUT™ Recombinant Ribonuclease Inhibitor (10777019, ThermoFisher Scientific), 200 U SuperScript™ IV Reverse Transcriptase (18090050, ThermoFisher Scientific), a final concentration of 5mM Dithiothreitol (DTT) (70726, ThermoFisher Scientific) and 1x SuperScript™ IV buffer in a total reaction volume of 20 µl. The thermocycler program for the second step was the following: 50 °C for 10 min, then 80 °C for 10min and 4 °C until further usage. The obtained cDNA was then 5x diluted with nuclease-free water to a final volume of 100 µl.

### Quantitative real-time PCR

The reaction mix per sample for quantitative real-rime PCR (qPCR) contained: 5 µl of the LightCycler 480 SYBR Green I Master Mix (04707516001, Roche), 2.5 µl cDNA and 2.5 µl primers in a total reaction volume of 10 ul. The reaction was performed in a LightCycler 480 (384) RT-PCR platform (Roche), using the LightCycler 480 Multiwell 384-well plates (04729749 001, Roche) and LC 480 Sealing Foil (04729757001, Roche). The program for qPCR was the following: 95 °C for 5 min; 45 cycles of (55 °C for 10 sec, 72 °C for 20 sec, 95 °C for 10 sec); melting curve (65-97 °C). The results were analyzed with the LightCycler 480 SW 1.5 software. Primers used for qPCR: RPS9 (QT00233989, Qiagen) as a reference gene, VIMP/SELS (QT00008169, Qiagen), IL2 (QT00015435, Qiagen), CSF2 (QT00000896, Qiagen), IL21 (QT00038612, Qiagen), CEBPG (QT00224357, Qiagen), E2F5 (QT00062965, Qiagen), IRX3 (QT00227934, Qiagen), RNF14 (QT00088291, Qiagen), ZBTB20 (QT00069776, Qiagen) and CTLA4 (QT01670550, Qiagen).

### Western blotting

Proteins were separated in Novex™ WedgeWell 4-20% Tris-Glycine Gels (XPO4202Box, Invitrogen), using the Novex™ Tris-Glycine SDS Running buffer (LC2675-4, Invitrogen). The proteins were transferred (dry transfer) using an iBlot2™ Gel Transfer Device (IB21001, Invitrogen) and iBlot2™ PVDF stacks (IB24002, Invitrogen). After the transfer the membranes were blocked in 5% milk in PBS with 0.2% Tween20 (PBS-T) for 1 hour at room temperature with gentle shaking before being incubated overnight at 4°C with the primary antibodies, diluted in 5% BSA in PBS-T with 0.025% sodium azide. The next day the membrane was washed three times for 10 min before and after incubation with secondary goat anti-rabbit HRP-coupled antibodies. The proteins were detected using the Amersham ECL Prime Western Blotting Detection Reagent (RPN2232, GE Healthcare Life Sciences) and visualized on the ECL Chemocam Imager (INTAS). If necessary, the contrast and brightness of the obtained whole gel pictures was adjusted using *ImageJ*. The signal intensity of the protein bands was quantified using *ImageJ* and normalized to that of the housekeeping gene GAPDH. For the quantification of phospho proteins, both the phospho and the total protein were normalized to GAPDH, before normalizing the phospho protein to the total protein.

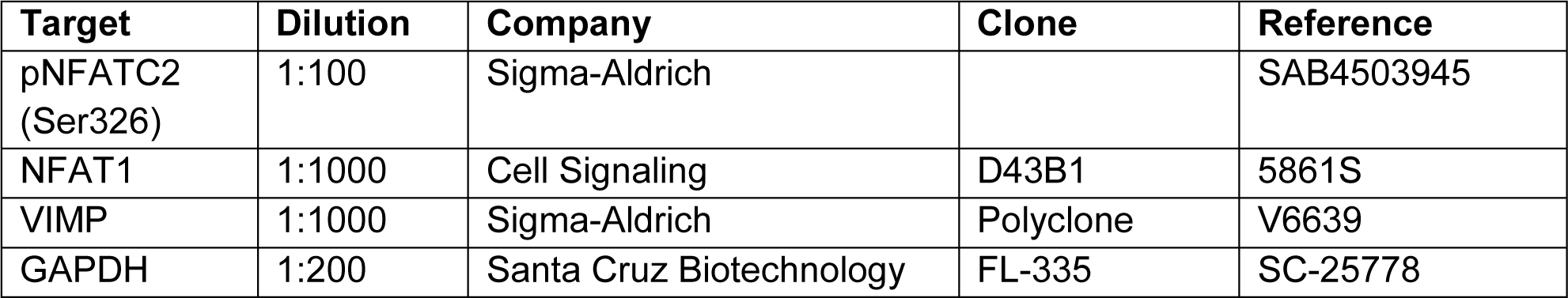

### Proliferation assay

The proliferation of the Teffs was assessed using the CellTrace™ CFSE cell proliferation kit (C34554, Invitrogen). The final concentration of 1 μM CFSE dye was used in our work. To label the cells, they were incubated for exactly 2 min and 45 seconds at RT in the dark. To stop the reaction, 10 ml FBS was added and the cells were centrifuged at 200 g for 10 min. After washing the cells in IMDM medium, the cells were subjected to the siRNA knockdown and counted. 10^5^ Teff in a 96-well plate were used for each condition and stimulated for 2 days with a ratio of 1:1 of irradiated Epstein Barr Virus (EBV) B cells as previously described (*29*). After the stimulation, the cells were stained for living cells using LIVE/DEAD® Fixable Near-IR Dead Cell Stain (L10119, ThermoFisher Scientific) (dilution 1:500) and acquired on a BD Fortessa™ analyzer. The data was analyzed in FlowJo 7.6.5.

### Cytokine measurement by Mesoscale discovery (MSD) platform

The cell supernatant was collected after centrifugation of the cells (250 g, 10 min) and the selected list of secreted cytokines (CSF2, IL2, IL21) was measured in the undiluted cell culture medium using the MSD U-PLEX Human Biomarker group 1 kit (MSD, K15067L-1) and following the manufacturer’s instructions. MESO QuickPlex SQ 120 instrument was used to read the plate and the data was analyzed with the MSD Workbench software.

### Cytokine measurement by Cytometric Bead Array (CBA)

The cell supernatant was collected after centrifugation of the cells and the secreted IL2 in the diluted cell culture medium (1:4 dilution) was measured using the IL2 Flex set cytometric bead array (CBA) (BD, 558270) following the manufacturer’s instructions. The acquisition was done on a BD Fortessa™ analyzer and the data was analyzed in FCAP Array™ v3.0.

### PhosFlow cytometry analysis

Following stimulation, the cells were immediately fixed by adding the same volume of prewarmed BD Cytofix Fixation Buffer (554655, BD) for 1h at 37 °C. After collecting the samples at all the different time points, they were then washed in FACS buffer and re-suspended in 200 µL of BD Phosflow Perm Buffer III (558050, BD) containing the antibodies for 30min at 4 °C. After washing the cells with FACS buffer, they were re-suspended in FACS buffer to be acquired on the BD Fortessa™.

The antibodies used are the following (**Table below**): VIMP/SELS (V6639, Sigma-Aldrich) (dilution 1:200) with Goat Anti-rabbit IgG H&L Alexa Fluor® 647 (A-21245, Invitrogen) (dilution 1:200), NFAT1 FITC (611060, BD) (dilution 1:50), phospho p38 MAPK (T180/Y182) Alexa Fluor 647 (562066, BD) (dilution 1:50), Anti-Human phospho NFATC1 (pS172) mAb (MAB5640, R&D Systems) (dilution 1:400), phospho NFATC2 (NFAT1) (S326) (SAB4503945, Sigma-Aldrich) (dilution 1:800), PE-Cy7 Mouse anti-ERK1/2 (pT202/pY204) (560116, BD) (dilution 1:50), phospho JNK1/2 (T182/Y185) (dilution 1:200) (558268, BD), phospho cJun (S63) (9261S, Cell Signaling) (dilution 1:200), phospho p105 NFκB1 (S933) (4806S, Cell Signaling) (dilution 1:400), phospho p100 NFκB2 (S866/870) (4810S, Cell Signaling) (dilution 1:400), phospho p65 (S529) (558422, BD) (dilution 1:50), phosphor RelB (S552) (4999S, Cell Signaling) (dilution 1:400), Anti-Rabbit IgG H&L Alexa Fluor 647 (ab 150079, Abcam) (dilution 1:1000), APC Goat Anti-mouse IgG (minimal X-reactivity) (405308, Biolegend) (dilution 1:200). For the acquisition a BD Fortessa™ was used and the data was analyzed in FlowJo 7.6.5.

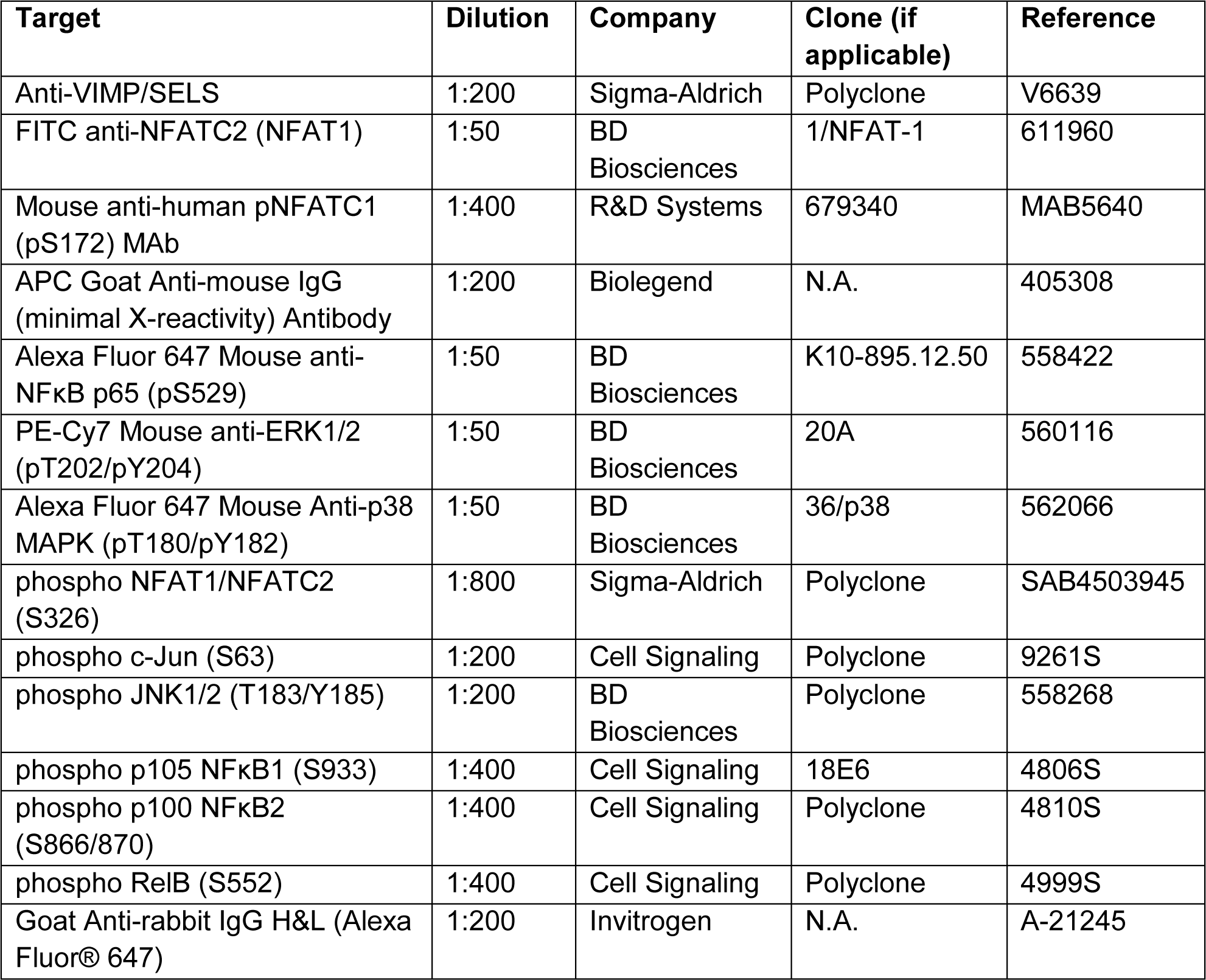

### Calcium/Ca2+ flux

To measure the calcium flux in Teffs, the cells were stained with mouse monoclonal [RPA-T4] anti-human CD4 FITC (555346, BD Biosciences) (dilution 1:100), LIVE/DEAD® Fixable Near-IR Dead Cell Stain (L10119, ThermoFisher Scientific) (dilution 1:500) and the calcium dye Indo-1 (I1203, ThermoFisher Scientific) (5 uM) for 60 min at 37 °C in supplemented IMDM medium as for the culture of Teffs. Following 3 washes with medium the cells were resuspended in 300ul of medium and incubated for another 15-30min at 37°C. The baseline of the calcium signal was measured for approximatively 30 seconds before adding the soluble CD3/CD28 antibodies (1:40) (10971, StemCell) or 100ng/ml Ionomycin (I0634, Sigma-Aldrich) to measure the activation-induced calcium flux. The cells were acquired on a BD Fortessa™ analyzer and the data was analyzed in FlowJo v10.5.

## Acknowledgement

We thank Annegrät Daujeumont, Alexandre Baron and Olga Kondratyeva for their expert technical supports. We particularly appreciate Luxembourg Red Cross for providing buffy coats to us.

## Author Contributions

C.C., N.Z., E.D. designed and performed experiments. C.C. analyzed the data and wrote the manuscript. S.F.R. performed parts of the experiments. R.B., M.O. and F.H. supervised the project. F.H. designed the project, oversaw the whole project and revised the manuscript.

## Statistical analysis

P values were calculated with paired two-tailed Student t test (Graphpad prism or Excel) as specified in Figure legend. All error bars represent the standard deviation.

## Data availability

Data availability: The microarray data have been deposited into Gene expression Omnibus (GEO) repository with the access code <u>GSE151266</u>. All other data and information needed to evaluate the conclusions of this work are presented in the Supplementary Figures.

## Supplementary Figures

**Supplementary Figure 1.**
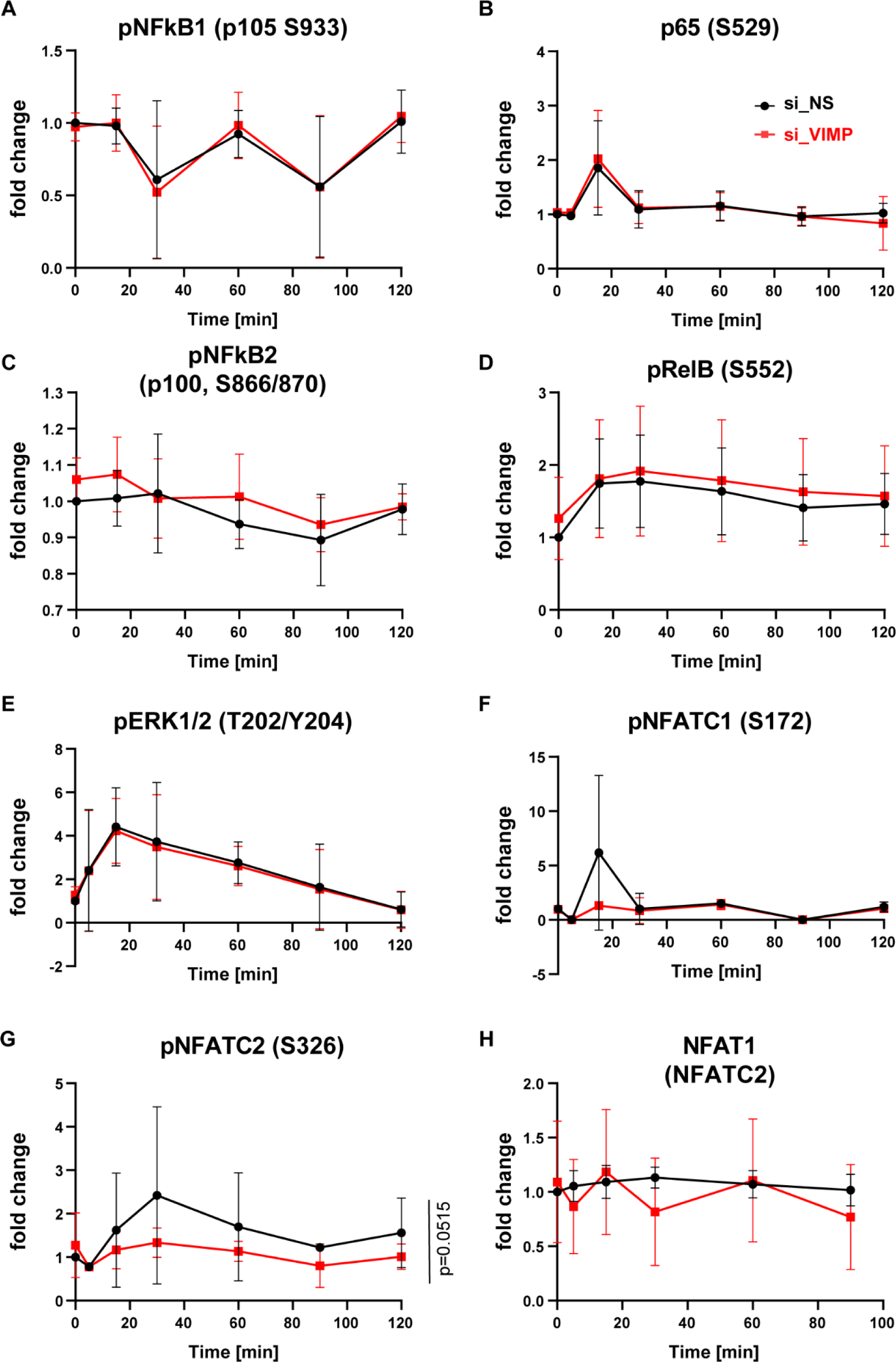
VIMP knockdown only affects the phosphorylation of NFATC2, not the other major signaling pathways downstream of the TCR. Phosphorylation of proteins involved in the major signaling pathways downstream of the TCR signaling in Teffs, assessed by flow cytometry at different time points following PMA/Ionomycin stimulation. Before stimulation, the cells were transfected with specific siRNA against VIMP (si_VIMP) versus non-specific siRNA (si_NS) for 1 day. (**G**) Only pNFATC2 was significantly decrease by VIMP knockdown. The other measured targets remain no significant change (**A**-**F, H**). The fold change was calculated by normalizing the geometric mean (Geomean) of the fluorescence intensities of all the conditions to that of the unstimulated control knockdown condition. Data are mean± standard deviation (s.d.). The P-values are determined by a two-tailed paired Student’s t test over time including the data at different time points. ns or unlabeled, non-significant; *P<=0.05, **P<=0.01 and ***P<=0.001. All the graphs represent the pooled flow cytometry data for the fold change from 2-7 independent donors.

**Supplementary Figure 2.**
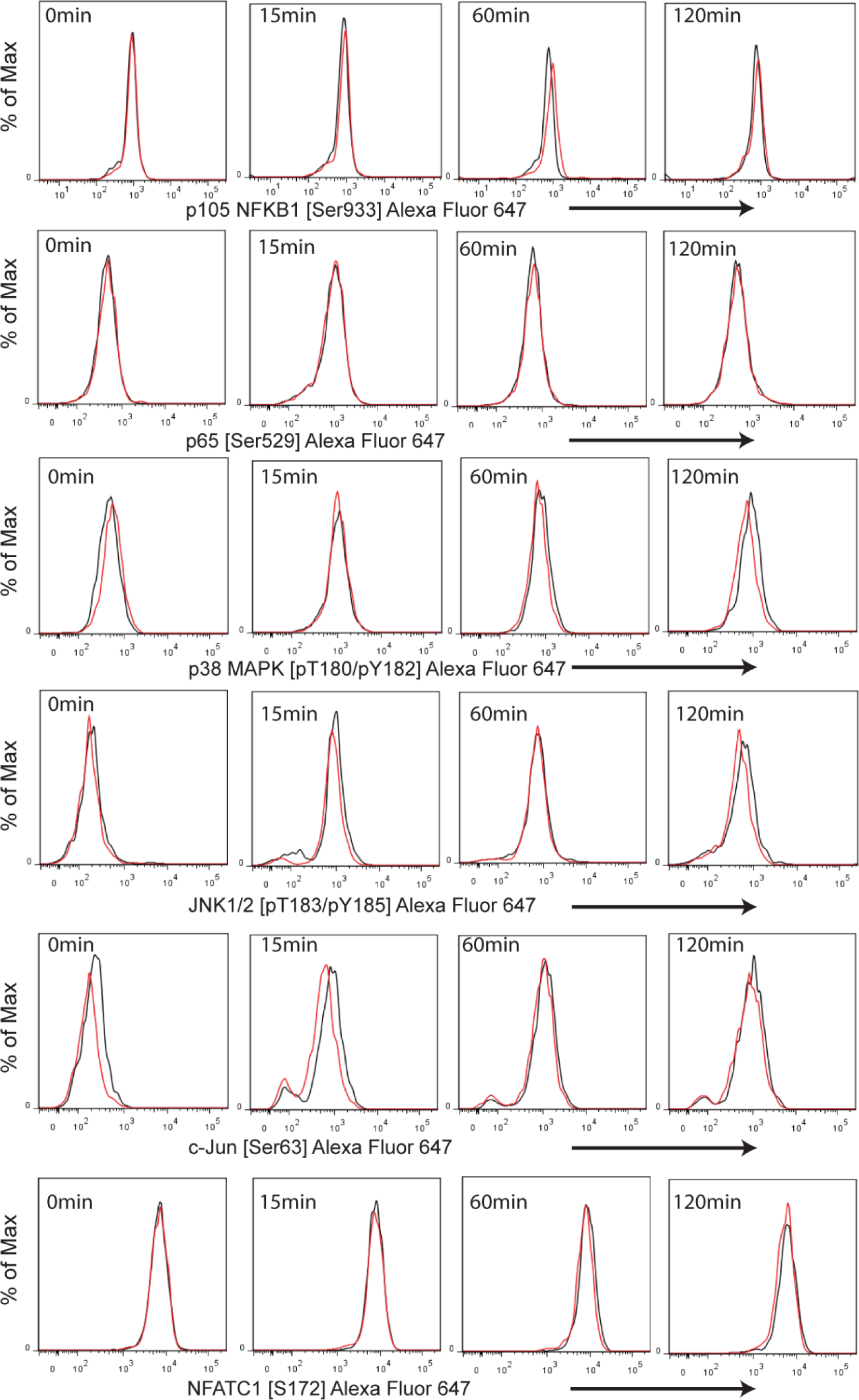
VIMP knockdown does not affect other major signaling pathways downstream of the TCR. Representative histogram overlay for the phosphorylation of major signaling transduction proteins downstream of the TCR signaling in Teffs, assessed by flow cytometry at different time points following PMA/Ionomycin stimulation. Before stimulation, the cells were first transfected with specific siRNA against VIMP (si_VIMP) versus non-specific siRNA (si_NS) for 1 day. No significant effects on the phosphorylation levels of MAPK (p38, ERK1/2, cJun, JNK1/2) pathways and canonical (p65, p105) or non-canonical (RELB, NFκB2) NFκB pathways during the first 120 min stimulation after siRNA knockdown in Teffs. The expression of total NFAT1 protein was also unaffected by VIMP knockdown. The numbers in x-axis indicate the geometric mean (Geomean) fluorescence intensity of the different proteins or phosphorylation sites. Data are mean± standard deviation (s.d.). The other measured targets remain no significant change (**A**-**G**). The P-values are determined by a two-tailed paired Student’s t test. ns or unlabeled, non-significant; *P<=0.05, **P<=0.01 and ***P<=0.001. All the graphs represent data from 2-7 independent donors.

